# Transient interactions drive the lateral clustering of cadherin-23 on membrane

**DOI:** 10.1101/2022.01.09.475209

**Authors:** Cheerneni S Srinivas, Gayathri S Singaraju, Veerpal Kaur, Sayan Das, Sanat K. Ghosh, Amin Sagar, Anuj Kumar, Tripta Bhatia, Sabyasachi Rakshit

## Abstract

Cis and trans-interactions among cadherins secure multicellularity. While the molecular structure of trans-interactions of cadherins is well understood, identifying the molecular cues that spread the cis-interactions two-dimensionally is still ongoing. Here, we report that transient, weak, multivalent, and spatially distributed hydrophobic interactions, requisite for liquid-liquid phase separations of biomolecules in solution, alone can drive the lateral-clustering of cadherin-23 on a membrane. No specific cis-dimer interactions are required for the lateral clustering. *In cellulo*, the cis-clustering accelerates the cell-cell adhesion and, thus, contributes in cell-adhesion kinetics along with strengthening the junction. Although the physiological connection of cis-clustering with rapid adhesion is yet to be explored, M2-macrophages that predominantly express cadherin-23 undergo fast attachments to circulatory tumor cells during metastasis.

## Introduction

Cadherins predominantly maneuver the active cell-adhesion processes for both vertebrates and invertebrates. Two modes of binding are known for cadherins, trans-binding, and cis-binding. While in trans-binding, the terminal extracellular (EC) domains of cadherins from opponent cells interact (Parisini et al., 2007, Haussinger et al., 2004), cis-binding is the lateral interactions among cadherins of same cell (Harrison et al., 2011). The trans-binding mode is well established from structural and kinetics studies in vitro as well as *in cellulo* localizations. The existence of cis-binding is inferred from the widespread localization of cadherins at cell-cell junction(Perez and Nelson, 2004) (Pertz et al., 1999). Unlike trans-dimers, direct existence of cis-dimers are not yet reported. One of the well-accepted proposals is that the trans mediates direct contacts among opposing cells via a diffusion (trap) kinetic approach and secures a cell-cell junction (2008; Troyanovsky et al., 2007). Approximately a cluster of five independent classical E(pithelial)-cadherin proteins diffuse across the membrane and form the trans-mediated cell-cell adherin junctions with neighboring cells (Wu et al., 2015). Lateral or cis-interactions, thereafter, influence the clustering of cadherins on the premature junctions and strengthen the association between cells. Two specific cis-interactions are proposed from the crystal lattice of N-, C-, or E-cadherins that extend the intercellular junction linearly (Wu et al., 2010, Shapiro and Weis, 2009, Boggon et al., 2002). What was lacking in the proposal is the molecular pathway that laid the transition from a linear array to the two-dimensional arrangement at the cadherin mediated cell-cell junctions. For type I cadherins, it is proposed that a combination of cis and trans-interactions via exclusive shape complementarity form the two-dimensional adherens surface (Wu et al., 2010, Wu et al., 2011). Another observation from the diffusion-dynamics of lateral clusters is the presence of nonspecific yet attractive interactions in addition to specific cis-binding that break the directional boundaries of interactions (Thompson et al., 2020). We propose that the transient and nonspecific intermolecular interactions can independently drive the two-dimensional arrangement of cadherins on cell membranes without any specific cis- or trans-interactions. Using a combination of biophysical and biochemical methods, we confirmed no existence of cis-dimers or higher-order oligomers of cadherins via lateral interactions.

Clustering of solute in a solution is a typical example of phase separation. In cell biology, such a liquid-liquid phase separation (LLPS) is common in the cytoplasm. It is the developmental origin of the membrane-less liquid compartments like nucleoli (Latonen, 2019), centrosomes(Mahen and Venkitaraman, 2012), Cajal bodies (Gall, 2003), and stress granules (Buchan and Parker, 2009). Relatively uncommon, but the existence of LLPS is also reported with proteins like Zonula Occludens (Beutel et al., 2019) and nephrin (Banjade and Rosen, 2014) that are anchored to cell-membrane and mediate multiprotein cell-adhesion, signal transduction. Transient, weak, and favorable nonspecific interactions among like-neighbors are thermodynamically responsible for phase separations of biomolecules. Our hypothesis is that such favorable interactions independently can drive cis-clustering of cadherins on two-dimensional confinement; without any trans- or stable cis-dimeric interactions.

We performed an unbiased *in silico* search using the catGRANULE algorithm (Mitchell et al., 2013, Klus et al., 2014) across the cadherin-superfamily of proteins and identified cadherin-23 (Cdh23) (NP_075859) protein with a high propensity to undergo LLPS. Cdh23 is one of the long non-classical classes of cadherins with 27 extracellular (EC) domains. The two outermost domains of Cdh23 from opposing cells interact homophilically in a trans-conformation and mediate strong cell-cell adhesion among tissues like the heart, kidney, muscle, and testis (Sannigrahi et al., 2019, Singaraju et al., 2019, Sotomayor et al., 2012). Similar to classical cadherins, the cellular junction of Cdh23 is continuous and engages in lateral clustering (Sannigrahi et al., 2019). Apart from cellular adhesions, Cdh23 also interacts heterophilically with protocadherin-15 (Pcdh15) in the neuroepithelial cells of the inner ear and serves as gating-springs for mechanotransduction in hearing (Sotomayor et al., 2010). Here we aimed to solve the molecular mechanism of cis-clustering of Cdh23 and their functional benefits. We performed a combination of photoinduced reactions and membrane biophysics to verify the existence of cis-dimers of Cdh23 in clusters. Modulating the clustering of Cdh23 using chemical spacers and blockers, we further deciphered the nature of interactions that drive the lateral clustering of cadherins. We confirmed that transient, weak, and nonspecific interactions independently drive the clustering. Hydrophobic association dominates such transient lateral interactions.

Free-standing cis-clustering of Cdh23 independent of trans-binding prompted us to decipher the functional role of clustering on cell adhesion. We measured a significant acceleration in the rate of cell-cell adhesions driven by cis-clustering. Disruption of cis-clusters by hydrophobic blockers drastically dropped the rate. Notably, while the toxicity of the phase-separated states has already been proposed for intrinsically disordered proteins(Elbaum-Garfinkle, 2019), the fast-aggregation of cells is a demonstration of the functional implication of LLPS in cell adhesion.

## Results

### Cdh23 exists as a single in solution

Cdh23 is an extensively elongated protein with an EC region of ∼315 kD. Traditional methods such as native gel electrophoresis and chromatography always failed to detect the higher-order cis-complexes for Cdh23 and many other classical cadherins (**Fig. 1 A**). The general argument is that weak binding-affinity is responsible for a ‘no-show’ of complexes in harsh experimental approaches. Here, we performed the photoinduced cross-linking of unmodified proteins (PICUP) to capture any stable or metastable dimers or oligomers of Cdh23 in the solution. PICUP is preferred for such identification due to its short reaction time, the ability to arrest short-lived assemblies, and most importantly, no alteration in the native proteins (Fancy and Kodadek, 1999). PICUP captures short-lived oligomers by forming radical-induced covalent bonds between interactomes. Followed by PICUP, we performed one-dimensional gel electrophoresis to obtain the stoichiometric distribution of dimer and monomer of proteins in equilibrium. To increase the detection ability to ≥ 3 ng /protein, we used silver staining of PAGE. We noticed a dimer band for Cdh23 in PICUP (**Fig. 1 B**). Reportedly, Cdh23 has a strong affinity to a trans-homodimer (Sannigrahi et al., 2019, Singaraju et al., 2019, Altin and Pagler, 1995). Upon blocking the trans-interactions using an excess of Cdh23 EC1-2 in solution, we detected no trace of Cdh23-dimers in the SDS-PAGE even for 5 μM of protein, indicating either no specific interactions for cis-dimerization or a significantly poor affinity of Cdh23 towards cis-dimerization (**Fig. 1 B**).

**Figure 1.**
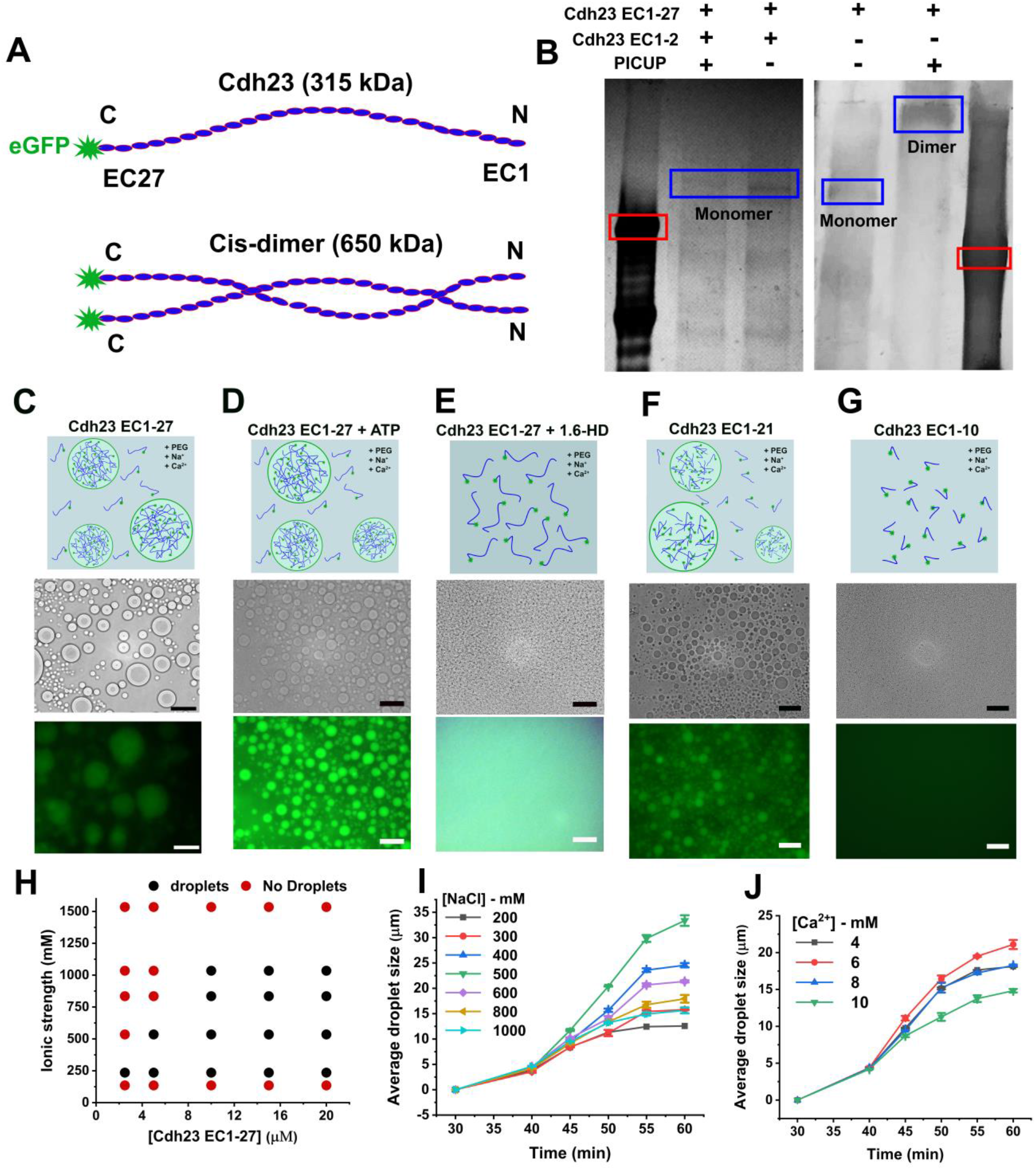
Weak, nonspecific, and transient interactions among EC domains facilitate the clustering of Cdh23 in the solution. **(A)** The schematic shows the monomer, and the hypothetical cis-dimer of Cdh23 EC1-27 recombinantly tagged with eGFP at C-terminus. **(B)** The silver staining of SDS PAGE gel shows the trans-dimer (∼650 kDa) of Cdh23 EC1-27 when subjected to PICUP. However, only a monomer of Cdh23 is detected on blocking the trans interactions of Cdh23 EC1-27. The trans homophilic interactions of Cdh23 EC1-27 are blocked by incubating with Cdh23 EC1-2. The rectangles (red) correspond to 315 kDa. **(C)** The schematic shows the assembly of biomacromolecules in solution via liquid-liquid phase separation. Representative bright-field (upper panel), and fluorescence (lower panel) images of liquid droplet-like condensates of Cdh23 EC1-27 at 60 min. Scale bar: 50 μm. **(D)** The representative bright-field and fluorescence images of liquid droplets of Cdh23 EC1-27 treated with ATP. Scale bar: 50 μm. (**E**) The schematic shows the dissolution of liquid droplets with the solution. The representative bright-field and fluorescence images of liquid droplets of Cdh23 EC1-27 in the presence of 1,6-HD. Scale bar: 50 μm. **(F)** The representative bright-field (upper panel), and fluorescence (lower panel) images of liquid droplets of Cdh23 EC1-21 at 60 min. **(G)** Representative bright-field (upper panel), and fluorescence (lower panel) images indicate the inability of Cdh23 EC1-10 to undergo LLPS. **(H)** Phase diagram of liquid droplets of Cdh23 EC1-27 relating protein concentration and ionic strength of the buffer with droplet formation. **(I)** Growth kinetics of liquid droplets (μm) at a varying concentration of NaCl. Error bars represent the standard error of the mean (SEM) with N=30 droplets. **(J)** The growth of liquid droplets (μm) of Cdh23 EC1-27 with time depends on Ca^2+^ concentration. Error bars represent the SEM for N=30 droplets.

### Cdh23 undergoes LLPS

Though Cdh23 does not form cis-dimers in solution, we observed a spontaneous phase separation (LLPS) of Cdh23 in the liquid phase from solution upon increasing entropy in the system. The fundamental difference in the nature of interactions in higher-order complexation and phase separation is the interaction strength and specificity. While protein-protein complex formation relies on relatively stable and specific interactomes, phase separation of a protein in a solution is driven by weak, spatially distributed, transient intermolecular interactions. To identify the EC regions in Cdh23 that are more prone towards phase separation, we performed an *in silico* search using the freely available catGRANULE algorithm. catGRANULE is a phase-separation prediction algorithm developed to estimate a propensity score (CDF) for proteins (of both mammalian and yeast) for LLPS on the basis of structural disorderness, length, and sequence of residues, and the content of arginine-glycine and phenylalanine-glycine. catGRANULE estimated a propensity score of 1.2 for the entire extracellular region (EC1-27) of Cdh23 (**Fig. S1**). Usually, a propensity score higher than 1 is considered as good LLPS candidate (Ambadipudi et al., 2017). We observed a spontaneous formation of liquid droplets of Cdh23 in the presence of inert crowder molecules like polyethylene glycol (PEG) above a critical protein concentration of 2.5 μM (**Fig. 1 C and H)**. The phase-separated state is stable over a range of ionic strength. To visually track the phase separations in real-time under a fluorescence microscope, we recombinantly tagged eGFP at the C-terminal of Cdh23 EC1-27 (**Fig. 1 A**). Fusion among floating droplets, the gold standard for liquid condensates, is also noticed among Cdh23 condensates, indicating the fluid nature of the droplets (**Video 1 A**). Further from the time-trace analysis of the fusion events, we measured an average fusion-time of 5.0 ± 1.2 s, in range with other proteins that undergo LLPS (Wang et al., 2019) (**Fig. S2** and **Video 1 B**).

Reportedly, Cdh23 mediates both homophilic and heterophilic trans-interactions. Two trans-interactions with two distinct binding affinities are reported between Cdh23 and Pcdh15 (Narui and Sotomayor, 2018). The most robust trans-conformation with a dissociation constant of < 1 μM was notified for the canonical variant. The second *trans*-conformation with a higher dissociation constant of 5 μM was observed for a truncated variant. It was, therefore, necessary to block the interference of the specific trans-interactions and measure the extent of droplet formation. Notably, the heterophilic trans-interaction with Pcdh15 has the highest affinity (**Table 1**) (Choudhary et al., 2020, Singaraju et al., 2019, Sotomayor et al., 2012). We accordingly blocked the trans-interacting sites of Cdh23 with ligand-protein Pcdh15 EC1-2. As a precaution, we first facilitated the heterophilic trans-interactions with an abundance of Pcdh15 EC1-2 (20 μM) in the experiment buffer and carefully altered the solution’s ionic strength from an unfavorable phase separation condition to a favorable state via dialysis (Materials and Methods). We observed LLPS of Cdh23 EC1-27 and Pcdh15 EC1-2 complex, indicating that the droplets are predominantly due to interactions among EC domains other than EC1-2 of Cdh23 (**Fig. S3**). Notably, the droplets of Cdh23 EC1-27 without Pcdh15 EC1-2 were more extensive than with Pcdh15 EC1-2, indicating that the additional trans-interactions contribute to the phase separation *in vitro* but are not essential for the LLPS. Overall, Cdh23 EC1-27 undergoes LLPS under physiological conditions, and the liquid droplets follow the characteristic feature of protein condensates.

**Table 1.**
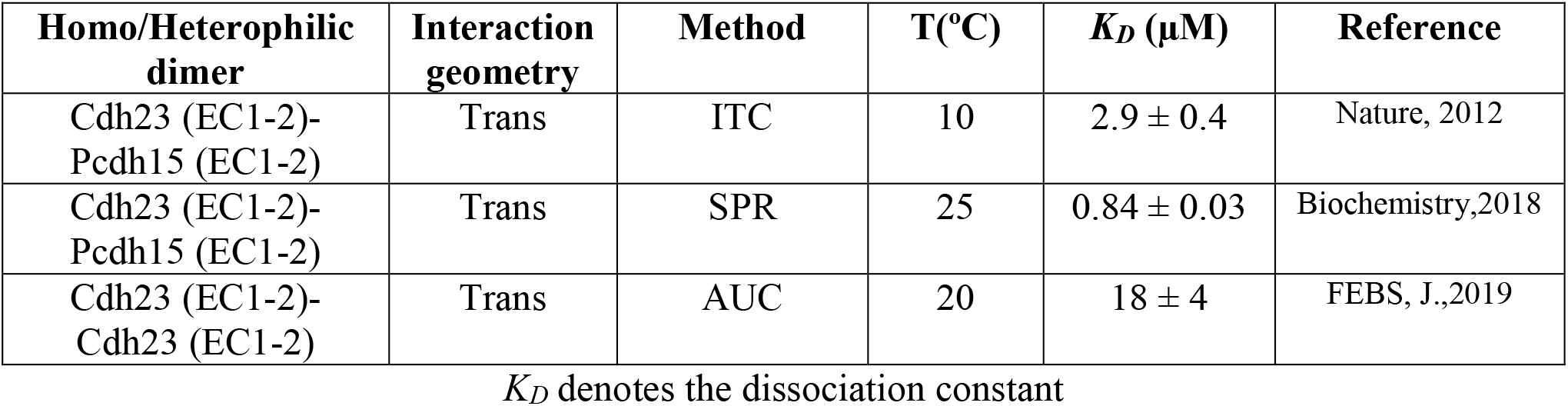
Binding-affinities of the heterophilic and homophilic trans-complexes of Cdh23 as reported.

| Homo/Heterophilic dimer | Interaction geometry | Method | T(°C) | $K_D$ (μM) | Reference |
| --- | --- | --- | --- | --- | --- |
| Cdh23 (EC1-2)-Pcdh15 (EC1-2) | Trans | ITC | 10 | $2.9 \pm 0.4$ | Nature, 2012 |
| Cdh23 (EC1-2)-Pcdh15 (EC1-2) | Trans | SPR | 25 | $0.84 \pm 0.03$ | Biochemistry, 2018 |
| Cdh23 (EC1-2)-Cdh23 (EC1-2) | Trans | AUC | 20 | $18 \pm 4$ | FEBS, J., 2019 |

$K_D$ denotes the dissociation constant

### Nature of interactions driving LLPS

What is the nature of interactions that facilitate the LLPS of Cdh23? Since the foundation of the dynamic condensed phase in a solution is transient and weak intermolecular interactions, the phase separated droplets rapidly and reversibly undergo mixing in the presence of interaction blockers. We used two different types of interaction blockers, Adenosine tri-phosphate (ATP) and 1,6-hexanediol (1,6-HD). ATP, being a condensed charge moiety, blocks the attractive electrostatic interactions for phase separation (Patel et al., 2017, Rice and Rosen, 2017, Hayes et al., 2018). In contrary, 1,6-HD, an aliphatic alcohol, is known to weaken the hydrophobic interactions critical for LLPS and inhibit the condensation of solutes to liquid droplets (Itoh et al., 2021, Duster et al., 2021). When we treated the liquid droplets of Cdh23 with ATP, we observed no change in the propensity for phase separation (**Fig. 1 D**). However, treatment of 1,6-HD disrupts liquid droplets of Cdh23 completely, indicating the predominant role of hydrophobic interactions in the LLPS (**Fig. 1 E**).

Among all EC domains, catGRANULE estimated a very high propensity of LLPS for EC10, 11, 14, and 16. EC2, 9, and 25 possessed a moderate propensity (**Fig. S1**). Residue-wise analysis indicated a major population of non-polar residues, including Val (V), Isoleucine (I), Leucine (L), Glycine (G), Proline (P), and Alanine (A) in the region. However, Cdh23 EC1-27 also possesses 440 negatively charged and 222 positively charged amino acid residues distributed throughout its length. It is thus inferred that the favorable interactions that drive the phase separation of Cdh23 are delicately balanced between repulsive electrostatic and attractive hydrophobic interactions. To experimentally delineate, we systematically varied the ionic strength of the buffer and estimated the optimal phase separation conditions. Ions screen the electrostatic repulsion among opposite charges and thus, favors hydrophobic interactions. Accordingly, we used monovalent and divalent ions (Na^+^ salts & Ca^2+^ salts, respectively) and monitored the droplet growth rate for optimization. We first varied Na^+^ ions keeping Ca^2+^ ions fixed at 4 mM. We noticed a gradual increase in droplet growth rate with increasing Na^+^ ions, reaching an optimum at 500 mM (**Fig. 1 I**). The phase separation of Cdh23 EC1-27 was noticed for 100 mM – 1 M of NaCl. Next, we set the Na^+^ ions to 500 mM, altered Ca^2+^ ions, and obtained an optimal phase separation at 6 mM of CaCl_2_ (**Fig. 1 J**). Overall, we observed that the screening of electrostatic charges did favor phase separation. Importantly, Na^+^ and Ca^2+^ ions beyond the salting-out range showed no LLPS for Cdh23 EC1-27.

To further substantiate the role of hydrophobic interactions in phase separation, we used two truncated variants of Cdh23, (i) Cdh23 EC1-21 containing domains with a high phase separation propensity of 1.1 and (ii) Cdh23 EC1-10 with low or no phase separation propensity of 0.8 (**Table 2**). Accordingly, we obtained spontaneous phase separation of Cdh23 EC1-21 in solution while Cdh23 EC1-10 did not undergo phase separation for a wide range of buffer conditions, including the requirements maintained for Cdh23 EC1-27 (**Fig. 1 F and G**).

**Table 2.**
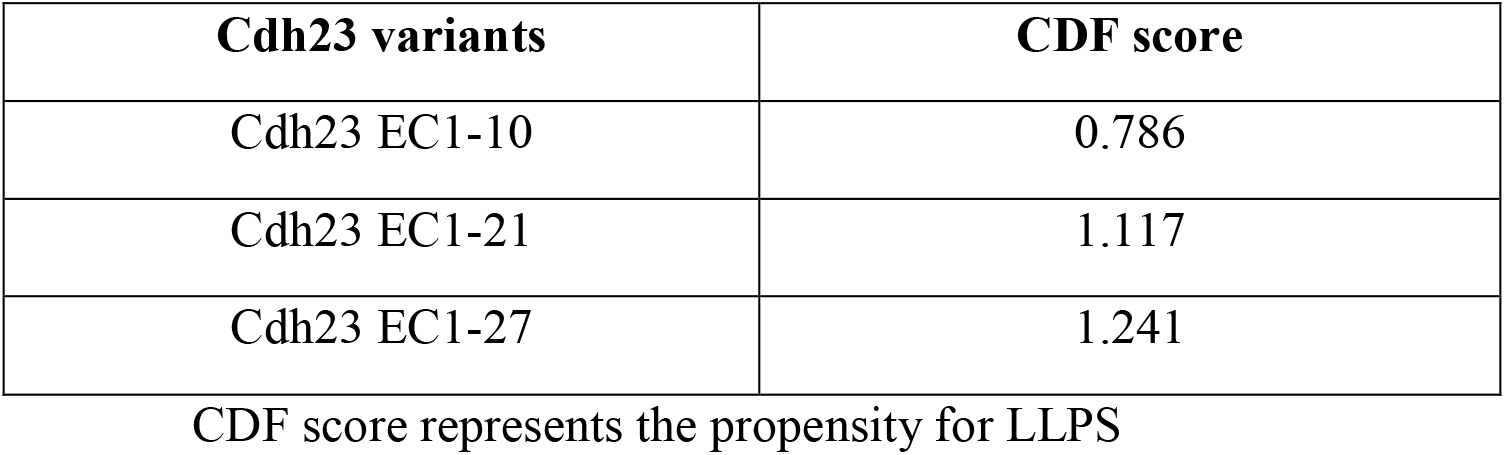
CDF scores estimated for Cdh23 variants from the catGRANULE algorithm.

### No cis-dimers of Cdh23 in the lateral clusters on lipid membrane

LLPS of biomolecules in a solution is the manifestation of weak and nonspecific transient intermolecular interactions. How effective is that transient interaction for lateral clustering on a membrane? Cadherins are trans-membrane proteins, and thus, the diffusion is restricted on two dimensions. Two-dimensional binding affinity is, therefore, significantly higher than the solution-based measurements. A dimensional restriction in diffusion and ordering in orientation enhances the binding affinity. It was thus speculative that the transient interactions can facilitate lateral clusters on a lipid membrane. To experimentally verify, we immobilized the eGFP modified Cdh23 on giant unilamellar vesicles (GUVs) and allowed free diffusion (See Methods). We noticed complete surface coverage of fluorescent proteins on GUVs. Fluorescence recovery after photobleaching (FRAP) has been a valuable tool for quantifying the clustering of molecules from the diffusion rates (Pincet et al., 2016, Kanaan et al., 2020). Accordingly, we performed FRAP experiments on GUVs where the C-terminal of Cdh23-GFP proteins was anchored via affinity (**Fig. 2 A, B and C**) (See Methods). We performed the FRAP experiments for all variants of Cdh23 and recorded the slowest diffusion-rate of 0.124 ± 0.005 μm^2^/s for Cdh23 EC1-21 and the fastest for Cdh23 EC1-10 is 0.60 ± 0.025 μm^2^/s (**Table 3**). The diffusion-rate of Cdh23 EC1-27 is 0.232 ± 0.007 μm^2^/s. The rate of diffusion is directly related to the cluster size. The slower the diffusion, the larger is the cluster size. Accordingly, Cdh23 EC1-21 forms a larger lateral cluster on the lipid membrane than the full-length Cdh23. This may not be surprising as CdhEC1-21 possesses a significant fraction of the long hydrophobic sequence with a high propensity for phase separation (**Fig. S1**). The cluster size for Cdh23 EC1-10 is the smallest, as estimated by catGRANULE.

**Table 3.** Diffusion coefficients measured for the clusters of Cdh23 variants (w/o 1,6-HD) anchored to GUV membranes.

| Cdh23 variants (w/o 1,6-HD) | Diffusion coefficient ($\mu\text{m}^2/\text{s}$)* |
| --- | --- |
| Cdh23 EC1-10 | $0.60 \pm 0.02$ |
| Cdh23 EC1-21 | $0.12 \pm 0.05$ |
| Cdh23 EC1-27 | $0.23 \pm 0.01$ |
| Cdh23 EC1-10 + 1,6-HD | $0.60 \pm 0.02$ |
| Cdh23 EC1-21 + 1,6-HD | $0.97 \pm 0.05$ |
| Cdh23 EC1-27 + 1,6-HD | $0.94 \pm 0.05$ |

\* It is important to note that the diffusion-coefficient of Cdh23 EC 1-27 and other smaller variants is closely dependent on the $[\text{Ca}^{2+}]$ ions in buffer.

**Figure 2.**
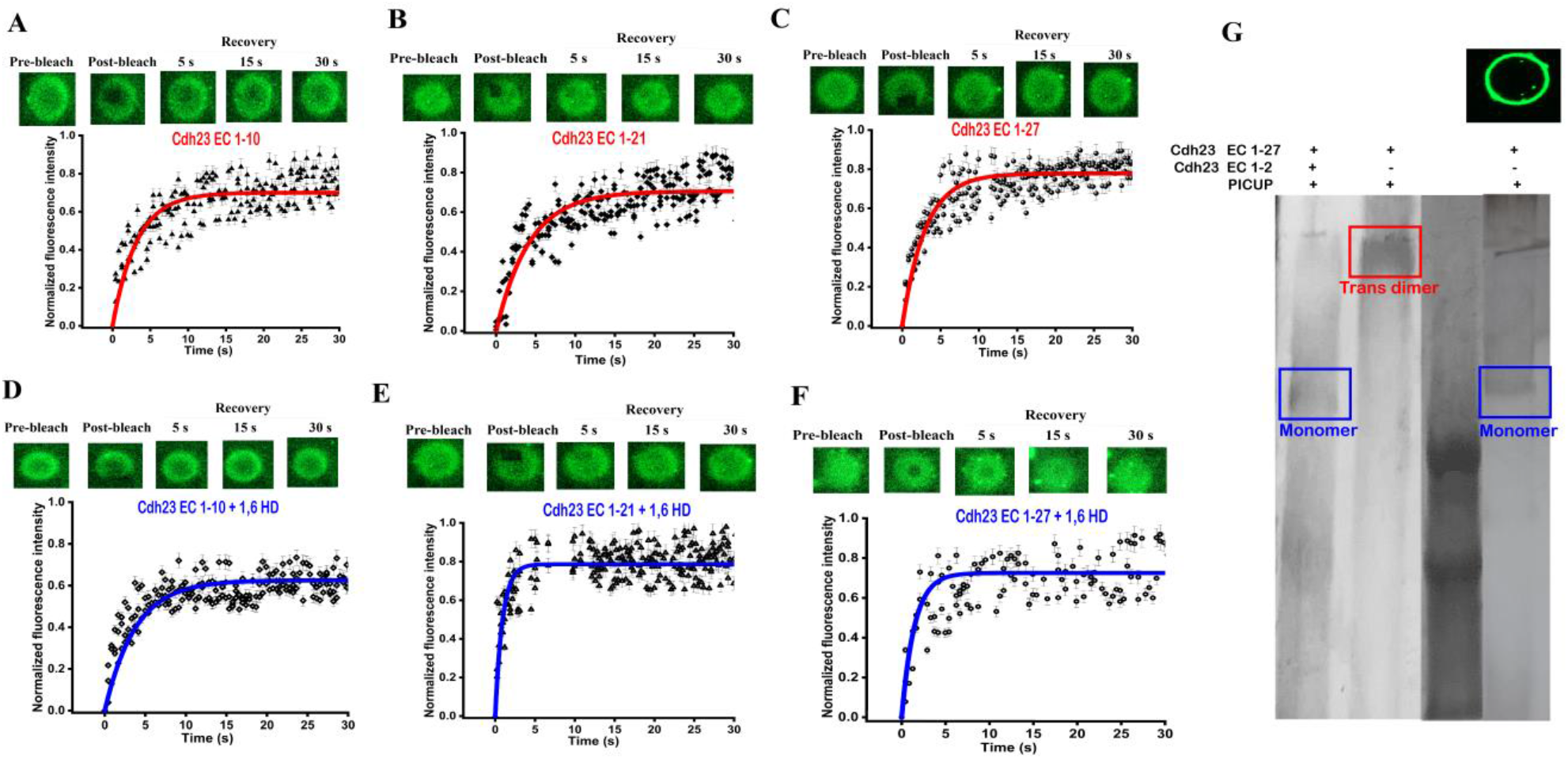
Cdh23 forms lateral clusters on GUV membranes. **A-F**. The fluorescence images of GUVs marking the pre-bleach, bleach, and post-bleach recovery regions at selected time points (upper panel), and the corresponding fluorescence recovery profile (normalized) (lower panel). **A, B, C** are for GUVs anchored with Cdh23 EC1-10, Cdh23 EC 1-21, and Cdh23 EC1-27, respectively. **D-F** are for the same GUVs anchored with Cdh23 EC1-10, Cdh23 EC 1-21, and Cdh23 EC1-27, respectively but after 1,6-HD treatment. The solid lines represent the fitting of the fluorescence recovery over time to the model (see methods). (**G**) PICUP on the clusters of Cdh23 EC1-27 on GUV membranes identifies only monomer band in the silver stained SDS-PAGE. Middle panel is the repeat of Figure 1B, indicating the relative position of the dimer band in red box. Upper inset shows the eGFP-fluorescent image of the representative GUVs used for PICUP.

Treatment of 1,6-HD that blocks the transient interactions enhances the rate of diffusion of proteins significantly to 0.97 ± 0.049 μm^2^/s and 0.94 ± 0.054 μm^2^/s for Cdh23 EC1-21 and Cdh23 EC1-27, respectively (**Fig. 2 D, E and F**). Noticeably, both variants diffuse at comparable rates in 1,6-HD. We measured, however, no change in the diffusion-rate for Cdh23 EC1-10 w/o 1,6-HD (**Table 3**). Importantly, 1,6 HD did not affect the membrane integrity as the diffusion dynamics of adsolubilized Nile Red dye remained unaltered w/o HD (**Table S1**). The treatment of 1,6 HD confirms that the weak, transient intermolecular interactions are predominant for lateral clusters of Cdh23 on a membrane. Moreover, we tracked the fluorescence recovery over a cross-section and noticed the recovery two-dimensionally, indicating the lateral clustering is distributed over the GUV surface without any influence of trans-interactions.

Does clustering of cadherins on lipid membrane facilitate specific interactions among neighbouring cadherins and generate cis-dimers or higher-order oligomers? To verify, we performed PICUP on lateral protein clusters on GUVs and monitored the existence of microstates from lateral interactions using SDS-PAGE of the PICUP samples. To our surprise, no higher-order bands appear in the PAGE, indicating the presence of no stable cis-dimers in the lateral cluster of Cdh23 on the lipid membrane (**Fig. 2 G**). Overall, our study with GUVs confirms that transient nonspecific interactions purely drive the lateral clustering of Cdh23, and no stable cis-dimers with specific interactions exist or produce in the lateral cluster. Arguably, a dimer of individual tip-link proteins is already reported using an electron micrograph (Kazmierczak et al., 2007). We also captured cis-dimer of Pcdh15 using PICUP on GUVs, however, no Cdh23 cis-dimer. Further, our observations on Cdh23 contradict the existence of specific cis-dimer interactions in the lateral cluster of E-cadherins on the lipid membrane.

### Lateral-clusters accelerate cell-cell aggregation

How does the lateral clustering of Cdh23 contribute to cells? Cadherins generally form anchoring junctions with the neighboring cells. Cdh23 is no different from the other family members and mediates vital cell-cell adhesion junctions in several tissues like the kidney, muscle, testes, and heart (Singaraju et al., 2019, Sannigrahi et al., 2019, Sotomayor et al., 2012, Li et al., 2019). We, therefore, verified the effect of the lateral condensations of Cdh23 in cell-cell adhesion, more importantly, where lateral-clustering precedes the trans-interactions. We hypothesized that lateral clustering of Cdh23 on a membrane will increase the effective intercellular interacting interface and accelerate cell-adhesion kinetics. We thus monitored the aggregation-kinetics of HEK293 cells exogenously expressing Cdh23 variants and estimated the adhesion-rate constants from the Von Bertalanffy kinetic model fit (West and Newton, 2019, Benzekry et al., 2014) (Materials and Methods). Generally, the complex tumor growth kinetics in *in vitro* spheroids and *in vivo* xenograft implants are quantitatively analyzed by fitting the data to Von Bertalanffy kinetic model that helps to forecast the future course of tumor progression. The time trace-tumor growth curves exhibit a simple curve pattern where the growth rate is not continuously exponential, i.e., their relative growth rate decrease with time (Benzekry et al., 2014). Our *in vitro* aggregation kinetics data also displayed a reduction in the growth rate over time. Moreover, the cell lines employed were cancerous. Hence, we considered the Von Bertalanffy kinetic model to quantify the cellular aggregations. Further, we measured the aggregation kinetics of the same cells but in the presence of 1,6-HD, where clustering of Cdh23 is blocked. As expected, we observed significantly rapid aggregations of HEK293 cells exogenously expressing Cdh23 EC1-27 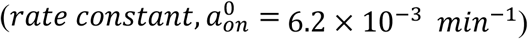 than the cells transfected with Cdh23 EC1-10 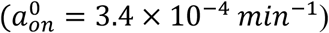 (**Fig. 3 A and B**) (**Table S2)**. We observed negligible aggregation with the untransfected cells. Treatment of 1,6-HD adversely affected the aggregation kinetics of HEK293 cells expressing Cdh23 EC1-27 (**Fig. 3 A and B**). Importantly, cells in all the experiments formed Cdh23 mediated matured cell-cell junction after incubation, warranting the unaltered functionality of Cdh23 in the presence of 1,6-HD. HEK293 cells treated with siRNA for endogenous Cdh23 showed no aggregation within the experiment time.

**Figure 3.**
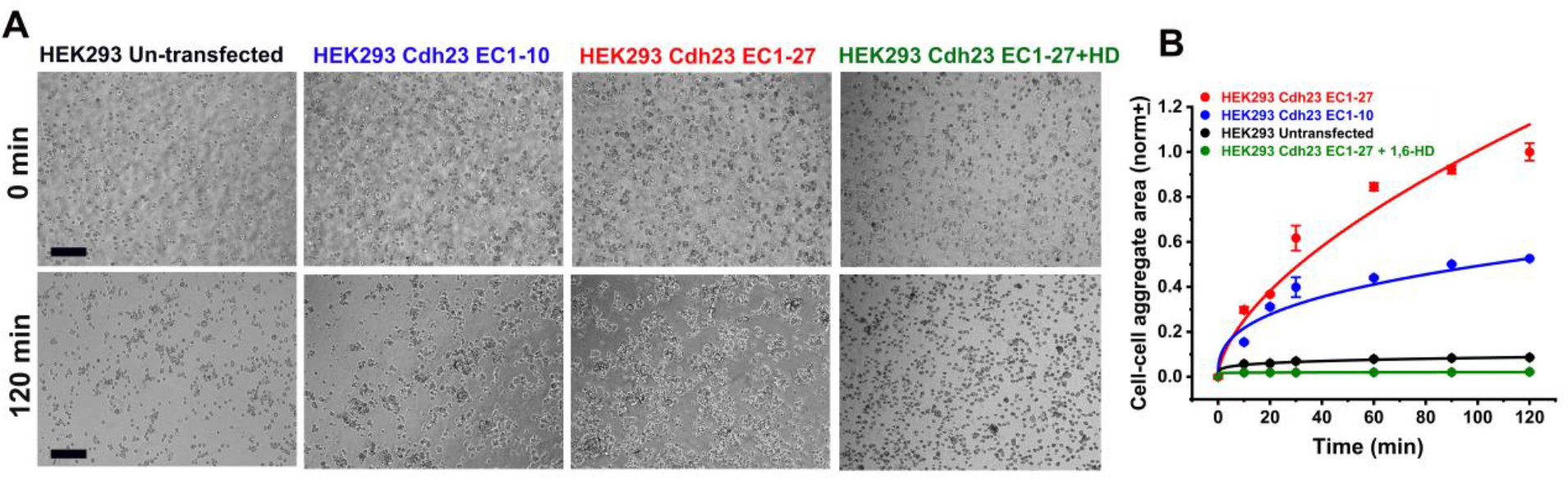
Lateral clustering of Cdh23 on membrane enhances the kinetics of cell-cell adhesion. **(A)** Time-stamp bright-field images of cell-cell aggregations of HEK293 cells untransfected (1^st^ column), transfected with Cdh23 EC1-10 (2^nd^ column), Cdh23 EC1-27 (3^rd^ column), and Cdh23 EC1-27 along with the treatment with 1,6-HD (4^th^ column). Scale bar: 50 μm. **(B)** The time-dependent growth of the cell-cell aggregation area of HEK293 cells exogenously expressing Cdh23 EC1-27 (red), Cdh23 EC1-27 and treated with 1,6-HD (olive), Cdh23 EC1-10 (blue), and untransfected cells (black). The error bars represent the standard error of the mean (SEM) with N=15 aggregates. The solid lines represent the fitting of aggregation kinetics to the Von Bertalanffy model.

Does cis-clustering on cell membrane depend on the extent of surface coverage by Cdh23? To check, we monitored the cell-aggregation kinetics among cancer cell lines, HEK293, HeLa, HaCaT, and A549 that express endogenous Cdh23 differentially. Our results from qRT-PCR and western blot, in corroboration with TCGA, indicate higher endogenous expression of Cdh23 in A549 and HaCaT cells and comparatively lower expression in HEK293 and HeLa cell-lines (**Fig. S4 A**). Amongst all in the list, HeLa has the least expression. We performed the cell-aggregation assays in the previously optimized buffer condition and noticed significantly faster cell-aggregations for A549 and HaCaT, than the low-expressing cell lines (HEK293 and HeLa). HeLa cells did not aggregate within the experiment time (**Fig. S5 A and B**) (**Table S1**). To conclude the differences in aggregation kinetics based on the extent of the lateral-clustering, we performed the aggregation kinetics of A549 and HEK293 cells in the presence of 1,6-HD. The aggregation kinetics of both cells dropped significantly in the presence of 1,6-HD and became comparable to HEK293 and HeLa cells (**Fig. 4 A and B**). Further, knock-out of Cdh23 in HEK293 and A549 cells using Cdh23-specific siRNA hindered the cell-aggregation significantly (**Fig. 4 A and B**). Overall, the kinetic data w/o 1,6-HD indicate that the differences in cell-adhesion kinetics of A549 and HEK293 cells are due to differences in the extent of lateral-clustering of Cdh23 (**Table S1**). When chemical treatments abolish the lateral-clustering, the individual Cdh23 molecules on the cell membrane follow the trans-mediated diffusion-trap kinetics for cell adhesion, similar to classical cadherins. While the lateral-clustering increases the effective binding interface on a cell membrane and kinetically facilitates the cell adhesion, the diffusion trap is instead driven by the binding affinities between partners.

**Figure 4.**
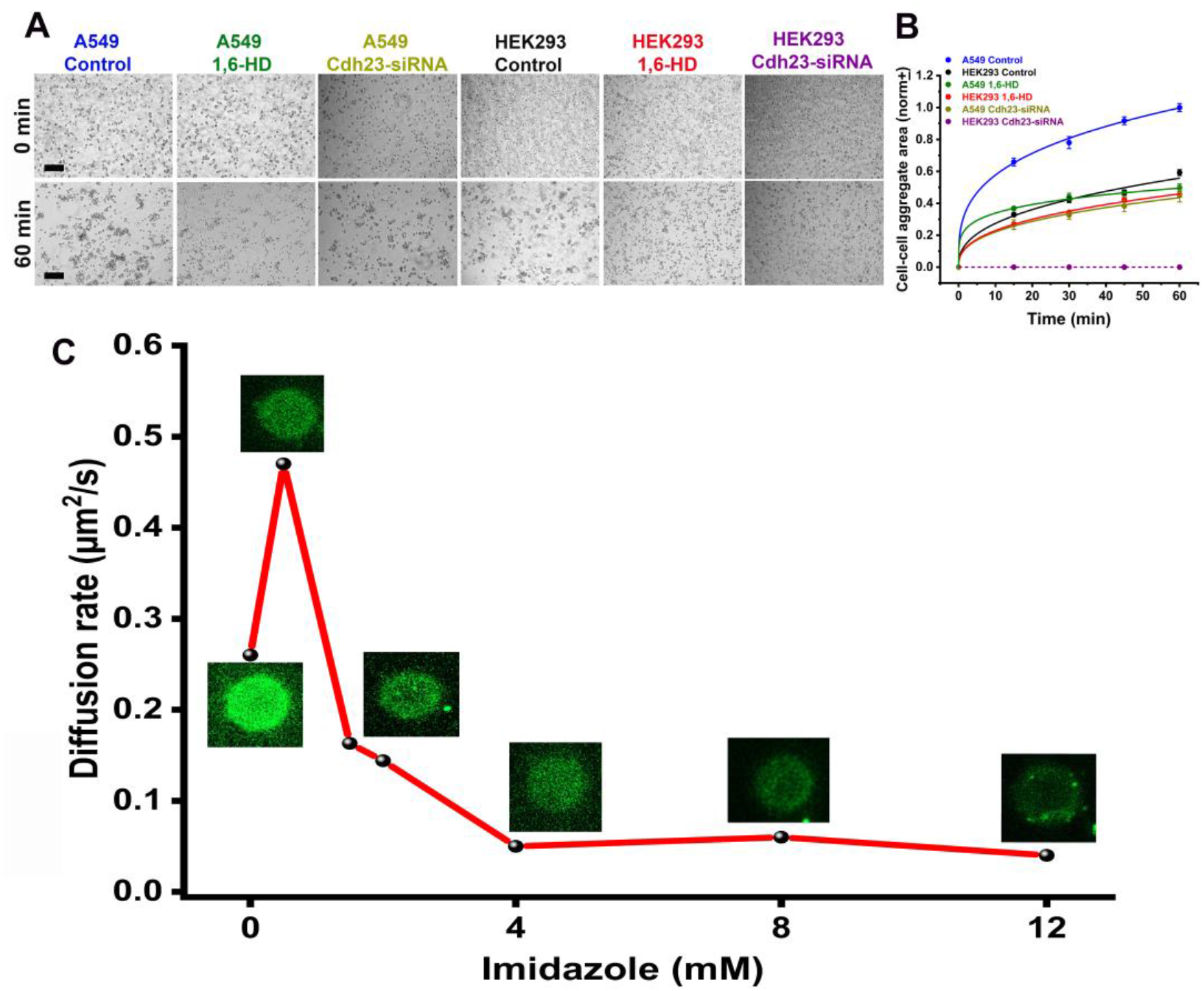
Effect of disrupted cis-clusters on cell-adhesion kinetics. **(A)** Time-stamp bright-field images of cell aggregates of A549 and HEK293 cells in the absence (control) and presence of 1,6-HD, and treated with Cdh23-siRNA. Scale bar: 50 μm. **(B)** Growth of cell-cell aggregation area (in μm^2^) with time for A549 cells in control (blue), treated with 1,6-HD (olive) and Cdh23-siRNA (dark yellow), and HEK2923 cells in control (black), treated with 1,6-HD (red) and Cdh23-siRNA (purple). Error bars represent the standard error of the mean (SEM) for N=15 aggregates. The solid lines represent the fitting of aggregation kinetics to the Von Bertalanffy model. **(C)** The gradual change in the diffusion-rates (μm^2^/s) of Cdh23 EC1-27 anchored to GUV membranes by His-tag with increasing concentration of imidazole (mM).

We further mimicked the differential extent of lateral-clustering of Cdh23 on GUVs. We titrated the Cdh23 EC1-27 attached GUVs with imidazole and monitored the change in clustering from the diffusion-rate. To note, the proteins are attached to GUV surfaces via Ni-NTA based affinity and treatment of imidazole replaces proteins from the GUV surfaces, thus reducing the extent of clustering. As expected, we first noticed a sharp rise in the diffusion-rate as we increased the imidazole from 0 M to 0.5 M (**Fig. 4 C**). However, upon further increasing imidazole in solution, the diffusion-rate dropped gradually. We noticed significant drop in green fluorescent signal from the GUVs due to the loss of proteins from the surfaces. While a rise in diffusion can be attributed by faster diffusion of relatively smaller clusters, a drop in diffusion-rate is a combining outcome of dilution and loss of clustering. A dilute surface inhibits lateral clustering and also allows proteins to travel long for fluorescence recovery. Importantly, treatment of imidazole did not alter the integrity of the GUVs as measured from the diffusion of adsolubilized Nile Red dye (**Table S1**).

### Fluidic nature of Cdh23 clusters on the cell membrane

To monitor the dynamics of Cdh23 at the cell-cell adhesion junction, we performed FRAP experiments on the intercellular junctions of HEK293 cells that are stably transfected with Cdh23. The protein was recombinantly tagged with eGFP at the C-terminal. We noticed localization of eGFP at the cell-cell junctions as expected for Cdh23 (**Fig. 5 A**) and photobleached a confocal volume. Next, we monitored the fluorescence recovery along a line across the photobleaching spot (**Fig. 5 B and C**). This is to identify if the recovery is from the new Cdh23 exported and recruited to the membrane by cells or diffusion of membrane-bound proteins. The fluorescence intensity profile across the line is expected to follow an inverted Gaussian profile with a deep at the center of the photobleaching spot (**Fig. 5 D**). We noticed a widening in the Gaussian profiles with recovery and characteristics of diffusion of active proteins from the surrounding membrane. Recruitment of new proteins, in general, recovers the fluorescence intensity without diluting the surroundings, thus with a little widening of the Gaussian width (Erami et al., 2016). Next, we plotted the width of the Gaussian (σ^2^) with time-lapsed after photobleaching and fit to the linear equation and estimated the diffusion-coefficient of Cdh23 clusters at the cell-cell junctions (**Fig. 5 E**) (Materials and Methods). The diffusion-coefficient of Cdh23 clusters at the cell-cell junctions is 0.6×10^−3^ + 0.1×10^−3^ μm^2^ s^-1^, 8-fold slower than the reported diffusion-coefficient of classical E-cadherin (D_eff_ = 4.8×10^−3^ + 0.3×10^−3^ μm^2^ s^-1^) clusters of ∼1000 molecules (Erami et al., 2016). Overall, our FRAP data indicates that Cdh23 at the cell-cell junction is fluidic and diffuse in clusters.

**Figure 5.**
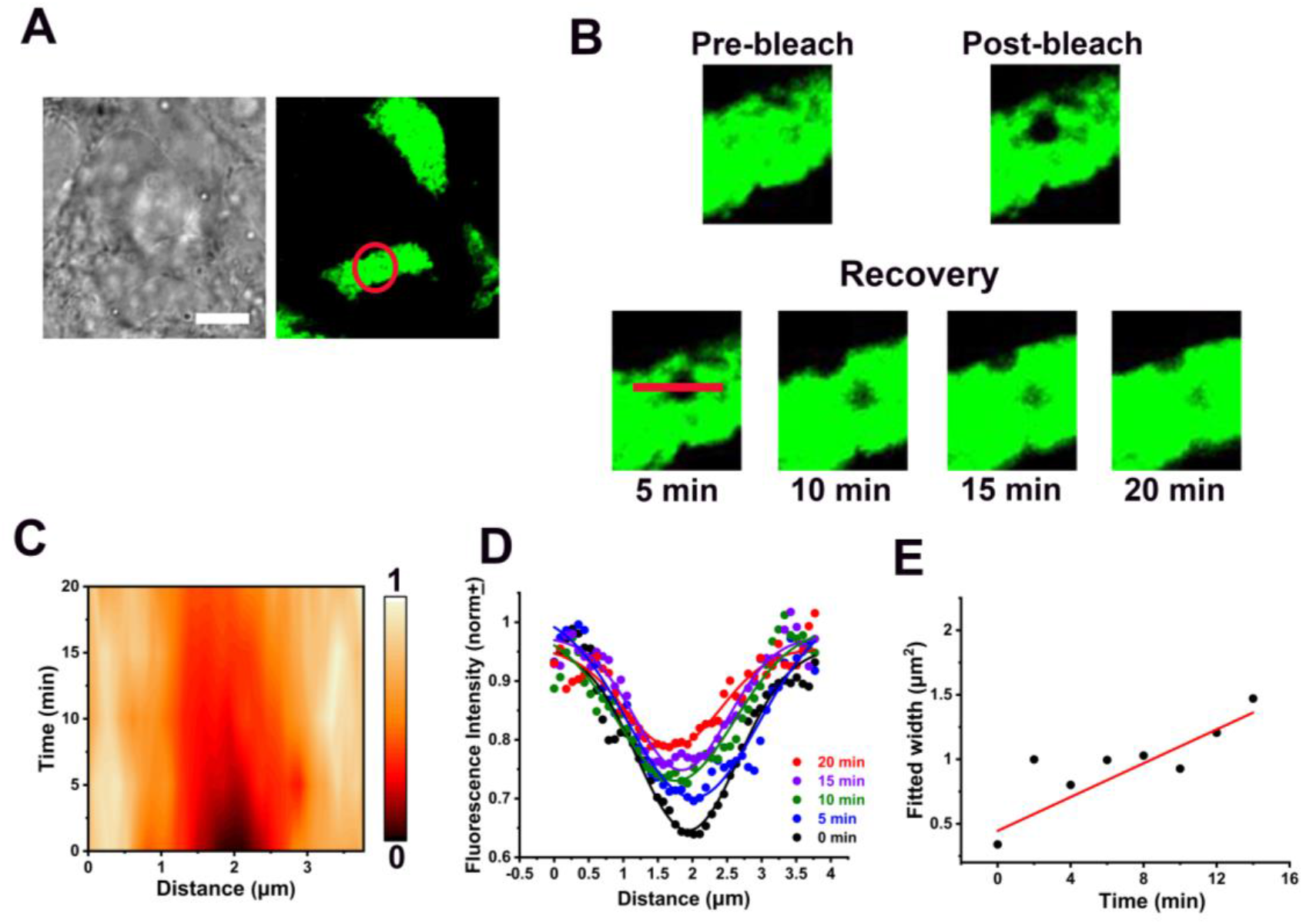
FRAP to probe the fluidic cis-clusters of Cdh23 at the cell-cell junctions. **(A)** Representative bright field and fluorescence images show the localization of Cdh23 at the cell-cell junction of HEK293 cells exogenously expressing Cdh23. Cdh23 is recombinantly tagged with eGFP at C-terminus. The circle (red) indicates the confocal volume for the FRAP experiment. Scale bar: 10 μm. **(B)** The cross-sections of fluorescence images indicate the regions of pre-bleach, bleach, and post-bleach recovery at the selected time points. The fluorescence recovery is monitored with time along the red line. The length of the line is 4 μm. **(C)** The contour plot represents the spatiotemporal distribution of Cdh23-eGFP along the red line. **(D)** The fluorescence intensity profile along the red line is plotted with the recovery time points. The solid lines are the Gaussian fits. **(E)** The widths (σ^2^) from the Gaussian fits are plotted with recovery time. The solid red line is the linear fit to the data to estimate the diffusion-coefficient. Error bars indicate the standard deviation obtained from the Gaussian fit.

## Discussion

At least two different molecular models are reported on the cis-clustering of classical cadherins. One of the models refers that the specific lateral intermolecular interactions among the EC domains drive the linear cis-clustering. The linear cis-interactions are, thereafter, expanded two-dimensionally at the junctions via intercellular trans-interactions (Wu et al., 2010). The other model, however, involves isotropic nonspecific interactions in addition to specific cis-dimer interactions to explain the two-dimensional spread of cis-clustering at the cell-cell junction (Thompson et al., 2020). It is, therefore, a long standing challenge in cell-cell adhesion to decipher the nature of interactions that spread a two-dimensional cell-cell junctions. We showed that weak, transient, nonspecific, and spatially distributed multivalent interactions that drive LLPS of molecules in a solution, can be responsible for the two-dimensional spread of Cdh23 clusters on a membrane even prior to any trans-interactions. The interactions are predominantly hydrophobic in nature.

Further, the cis-clusters of cadherins are primarily known to strengthen the trans-interactions at the cell-cell junction (Wu et al., 2010). Our experimental observations, however, identify the independent role of lateral clustering on cell-adhesion kinetics. We noticed that Cdh23 could condensate to clusters at the cell membrane independent of trans-mediated anchorage to neighbouring cells and as super-adhesive, can rapidly seize the floating cells into aggregates. Though the implication of rapid cell-cell adhesion/communication in physiology or life sciences is not yet clear. Cdh23, among other cadherins, is significantly overexpressed in tumor-infiltrating M2-type macrophages (Poczobutt et al., 2016) and microglia (Zhang et al., 2016, Zhang et al., 2014) (**Fig. S4 B**). M2-type macrophages associate with the circulatory tumor cells (CTCs) on the go and help in metastasis. A quick cell-cell adhesion is thus essential in this process. Though speculative, the fast adhesion between M2 macrophages and CTCs is facilitated by the condensed Cdh23 droplets.

Moreover, Cdh23 may not be the only cadherin in the family that may undergo lateral clustering on membrane via weak transient interactions that drive LLPS of macromolecules in a solution. The catGRANULE algorithm estimated the CDF score of more than 1 for many other cadherins (**Table S3**), considering only the ectodomains. In general, with a CDF of more than 1, the proteins tend to phase separate via weak and transient intermolecular interactions. Accordingly, Fat-cadherin, Dacshous-cadherin, desmosomes, and cadherin-22 have CDF scores of more than 1 and can undergo LLPS. Interestingly, most of these cadherins are associated with special cell-cell junctions. For instance, fat-cadherin and dacshous cadherin-mediated heterophilic junctions exclusively regulate the epithelial cell-size dynamics (ECD) under the mechanical cues during morphogenesis (Kumar et al., 2020), desmoglein-2 forms heterophilic interactions with other isoforms of desmosomal cadherins and form Ca^2+^-independent hyper-adhesive desmosomal junctions in tissues like skin, heart that are exposed to physical forces.

## Conclusion

The distinctive features of nonspecific and transient lateral intermolecular interactions, stretchable and tunable to different sizes and shapes, may be helpful in cell-cell junction, which routinely experiences mechanical assault. Our results address the molecular cues that spread the lateral clustering of cadherin-junction in two-dimensions and decipher the kinetic control of cadherin junctions for the first time. Identifying the physiological or pathological phenomena that strongly depend on the rapid cell-cell communication and adhesion may open up another exciting field in cell-adhesion.

## Materials and Methods

### Cloning of domain deletion mutants of Cdh23

The full-length Cdh23 (NP_075859), consisting of 27 EC domains, a transmembrane domain, and a cytoplasmic domain, was a generous gift from Dr. Raj Ladher, NCBS, Bangalore. Using this construct, we recombinantly generated domain deletion mutants. We have subcloned the same construct in pcDNA3.1 (+) plasmid, which codes for Neomycin resistance. All the constructs were cloned between NheI and XhoI restriction sites with (S)-Sortase-tag (LPETGG)-(G)-eGFP-tag and (H)-His-tag; SGH-tag at downstream (C-terminus of the protein) in the same order. All the recombinant constructs were verified through double digestion, PCR amplification, and DNA sequencing.

### Protein expression and purification

All recombinant Cdh23 variants for *in-vitro* studies were expressed in the ExpiCHO suspension cell system (A29129 ThermoFisher Scientific), following the prescribed protocol for transfection in ExpiCHO cells. After seven days, the culture media was collected by pelleting down the cells at 2000 rpm for 15 min at room temperature. The media was then extensively dialyzed against the dialysis buffer for 48 hours and intermittently changed the buffer every 8 hours. The dialyzed media with proteins were purified using affinity chromatography using Ni-NTA columns. The purity of the samples was checked using SDS-PAGE. Finally, the presence of protein was confirmed using western blotting with specific antibodies against GFP, Cadherin-23, and his-tag.

### *In vitro* droplet formation assay

All purified proteins were prepared in a buffer containing 20 mM HEPES, pH 7.5, 100 mM NaCl, and 4 mM CaCl_2_. Before each experiment, the proteins were centrifuged at 15000 rpm at 4°C for 10 min to remove possible nonspecific aggregates. Then proteins were adjusted to reach designated concentrations. Each protein mixture (14 μM for each component) was injected into a homemade chamber and imaged using a Leica microscope (Leica DMi8) using a 40X objective lens. The time-lapse images were taken under bright-field and fluorescence filters. All the assayed droplets were thicker than 6 μm in height, so the central layers of optical sections were chosen for quantification. Over 10 or more droplets were measured for each protein to generate the phase diagram of the condensed phase. The images were analyzed by ImageJ, and the quantification was performed by Origin software.

### PICUP

About 5 μM of purified Cdh23 was added with a metal complex containing 15 mM sodium phosphate, pH=7.5, 150 mM NaCl, 0.125 mM [Ru(bpy)_3_Cl_2_], and an electron acceptor, 2.5 mM of Ammonium persulfate (APS), followed by the irradiation of visible light for 0.5 sec. With no delay, the reaction mixture was quenched with 7 μl of 4X SDS-gel loading dye and heated at 95 ºC for 5 minutes. Next, SDS-PAGE and silver-staining were performed consecutively to visualize the cross-linked product. Cdh23 EC1-2 (5 μM) was added to block the trans-homodimerization of Cdh23 during PICUP experiments.

### GUVs preparation

The mixture of DOPC (850375P, Sigma-Aldrich), DGS NTA Ni (790404C, Sigma-Aldrich), and DSPE-PEG Biotin2000 (Avanti) lipid solutions dissolved in chloroform was used to prepare GUVs. We followed the electroformation method to prepare the GUVs (Bhatia et al., 2017). Briefly, the calculated amounts of lipids mixture (1 mM DOPC: 1 mM DGS NTA Ni) were spread on the plates and allowed the solvent to evaporate under vacuum. GUVs were harvested in a 350 mM sucrose solution. The formation of GUVs was visualized using phase contrast and fluorescence microscopy (Leica, Dmi8), where the GUVs were labeled with Nile red (1 μM), a fluorescent dye (72485, Sigma-Aldrich).

### Tethering GUVs to the functionalized surface and the specific attachment of the protein to GUV membranes

Glass base petri dishes were extensively cleaned with piranha and KOH etching before the protein incubations. The glass surfaces were coated with 1 mg/mL of BSA protein (Albumin, biotin-labeled bovine, A8549 Sigma-Aldrich) by incubating for 2 hours, followed by a wash with MilliQ water. Next, the surfaces were incubated with 0.1 mg/ml of Streptavidin stock (S4762, Sigma-Aldrich) for 1 hour and then washed with water. The GUVs were incubated on the modified surface for 4 hours to anchor them to the surface through the non-covalent interactions between biotin-streptavidin. The surface adhered GUVs were incubated with ∼ 6 μM of GFP-tagged proteins at 4 ºC for overnight.

### FRAP and analysis

FRAP experiments were performed at the polar region of the tethered GUVs using a super-resolution microscope (ZEISS LSM 980 Airyscan 2) at 63X magnification. The experiments were performed in the presence and absence of 1,6 hexanediol in the buffer.

The fluorescence intensity (normalized) at the selected region was measured for three independent experiments and plotted against the time. Recovery half lifetime (τ_1/2_) was estimated by the global fit of fluorescence intensity profile with recovery time using the following equation, *f*(t) = A(1 − e^-τ t^), where A indicates the mobile fraction of proteins and 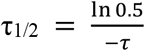

The diffusion coefficient for Cdh23 localized on vesicle membranes was calculated using the equation 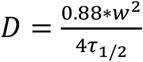, where W is the radius of the region selected to study FRAP.

### LLPS disruption studies

To disrupt the liquid-like clusters, the cells were incubated with 2% (w/v) of 1,6-Hexanediol (H11807, Sigma-Aldrich) for 1 hour. Similarly, 10% (w/v) of 1,6-Hexanediol was used to disrupt the liquid droplets of Cdh23 EC1-27 *in-vitro*.

### Western blot and qRT-PCR

The adherent cancer cell lines HeLa, HEK293, A549, and HaCaT were obtained from NCCS, Pune. All cells were cultured in high glucose DMEM media (D1152, Sigma-Aldrich) containing 10% FBS and 5% CO_2_. We followed the standard protocol(Hirano, 2012) for the western blotting of the lysates from the mentioned cell lines. Cadherin-23 (HPA017232, Sigma-Aldrich and PA5-43398, Invitrogen), eGFP (A11122, Invitrogen), and His-tag (11965085001, Roche) antibodies were used to detect the proteins.

RNA from different cancer cell lines was extracted using RNA isolation kit (Bio Rad) and treated with DNAse using DNAse 1 kit (AMPD1, Sigma-Aldrich). cDNA synthesis was done using a cDNA synthesis kit (Bio Rad). qRT-PCR was performed with the primers probing Cdh23 using the real-time PCR system (CFX96 Bio Rad).

### Cell-aggregation assay

After 30 hours of post-transfection, the cells were washed gently with PBS and then resuspended in Hank’s buffer supplemented with 10mM Ca^2+^ ions to a final cell count of 10^5^ cells. Hank’s buffer behaves like an incomplete media maintaining the osmolarity of the solution with the cells avoiding any bursting or shrinking of cells throughout the entire duration of the assay. After resuspending, the cells were imaged with a bright-field filter at 10X magnification using a Leica Inverted Microscope (Leica DMi8) over a time trace for 2 hours. The images were collected at 10 min, 15min, 30 min, 45 min, 60 min, and 120 min when all the cells aggregated completely. The cell-aggregates at 5 different surfaces were considered at each time point, and the sizes were measured from the bright field images using ImageJ software (NIH, Bethesda). The image analysis for measuring the area of each aggregate was done in ImageJ software. The aggregates with at least 5 cells are considered for the analysis. The mean area of aggregates over five different focal positions was measured and plotted against time. The aggregate size was compared over varying domain lengths for Cdh23. We performed all cell aggregation experiments at fixed cell numbers. 1% (w/v) of 1,6-Hexanediol was added to Hank’s buffer to disrupt the LLPS during the cell-aggregation assays.

### Fitting the cell aggregation data to a model

We have used the Von Bertalanffy model (West and Newton, 2019, Benzekry et al., 2014) to quantify the rate of cell-cell aggregation for HEK293 cells transfected with Cdh23 variants and cell lines differentially expressing Cdh23. The net rate of cell aggregation is proportional to the total area of aggregate. In the absence of any dissociation in our experiment timescale, we have neglected the loss term in the equation (a special case of the Von Bertalanffy model). Finally, we fit the cell aggregation data for an experimental condition over time using the following rate-equation:

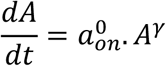

The model is solved and written explicitly as, 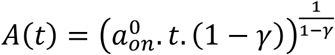, where, 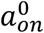 represents the rate-constant, *A* represents the area of aggregate, *t* is the independent variable (time), and *γ* represents the growth of aggregate. 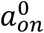 is an inherent property and *γ* is dependent on the cell types and their heterogeneity.

### Live cell imaging and FRAP analysis

Stably expressing Cdh23 HEK293 cells grown for confluency on a 35mm glass-base petri dish were used for imaging. A super-resolution microscope (Zeiss LSM980 Airyscan 2) was used to image the cells maintained at 37 ºC and 5% CO2. FRAP was performed on a confocal volume of 1 μm diameter at the cell-cell junctions where localization of eGFP was noticed. ImageJ software was used to measure the fluorescence intensity profiles of the line segment of 4 μm drawn across the photobleached region (line-scan analysis). The fluorescence intensity profiles (normalized) at different time points were fit to the Gaussian function in origin software. The fitted widths obtained at different time points were plotted against recovery time, and fit to linear regression to estimate the diffusion coefficient from the slope (Erami et al., 2016).

## Acknowledgments

This work was supported by the DBT/Wellcome Trust India Alliance Fellowship [grant number: IA/I/15/1/501817] awarded to SR.

SR acknowledges and thanks Dr. Raj K. Ladher, NCBS, for generously donating the Cadherin-23 plasmid. SR acknowledges the financial support by the DBT/Wellcome Trust India Alliance, Indian Institute of Science Education and Research Mohali (IISERM), and Centre for Protein Science Design and Engineering (CPSDE), Indian Institute of Science Education and Research Mohali. GSS, CSS, VK, and AS sincerely thank IISERM for financial support. SD thanks DBT/Wellcome Trust India Alliance for Project Assistant Fellowship awarded to SR. AK is grateful to DST Inspire for funding.

We acknowledge department of biological sciences, IISER Mohali, for providing the super resolution microscope facility granted by SERB for the Fund for Improvement of S&T Infrastructure in Universities and Higher Educational Institutes (FIST) Program.

We thank Mr. Taseen Ahmad for assisting in acquiring super-resolution microscopy data and Mr. Hemraj for helping us with the preparation of GUVs.

The authors declare no competing financial interests.

## Author Contributions

SR has supervised the project. GSS, SD, AS, and AK did the cloning, expression, and purification. GSS and CSS recorded and analysed droplet condensation and cell-aggregation experiments. CSS carried out the curve fitting on cell-aggregation. CSS did the live-cell imaging and FRAP experiments. TB, VK, and SR has designed the GUV experiments. TB has supervised the GUV experiments and the corresponding data-analysis. VK performed PICUP experiments. VK, SKG, and TB performed GUV experiments and analysed FRAP on GUVs. GSS, VK, and CSS made the figures. SR and CSS wrote the manuscript. GSS, CSS, VK, and SR edited the manuscript.

CSS, GSS, and VK have contributed equally in this work.

## Abbreviations

AUC: Analytical UltraCentrifugation Cdh23 Cadherin-23
CDF: Cumulative Distribution Function CD Cytosolic domain
EC: Extracellular
EC1-27: Extracellular 1-27 domains
eGFP: Enhanced Green fluorescent protein
FRAP: Fluorescence Recovery After Photobleaching
GUV: Giant unilamellar vesicles
1,6-HD: 1,6-Hexanediol
ITC: Isothermal titration calorimetry
LLPS: Liquid-liquid Phase Separation
Pcdh15: Protocadherin-15
PICUP: Photoinduced cross-linking of unmodified proteins
qRT-PCR: Quantitative Real-time polymerase chain reaction
SPR: Surface Plasmon Resonance
TM: Transmembrane

