## Supplementary File for "Transient interactions drive the lateral clustering of cadherin-23 on membrane"

**This file includes:**

**Figures S1 to S5**

**Legends for videos 1 A and B**

**Tables S1 to S3**

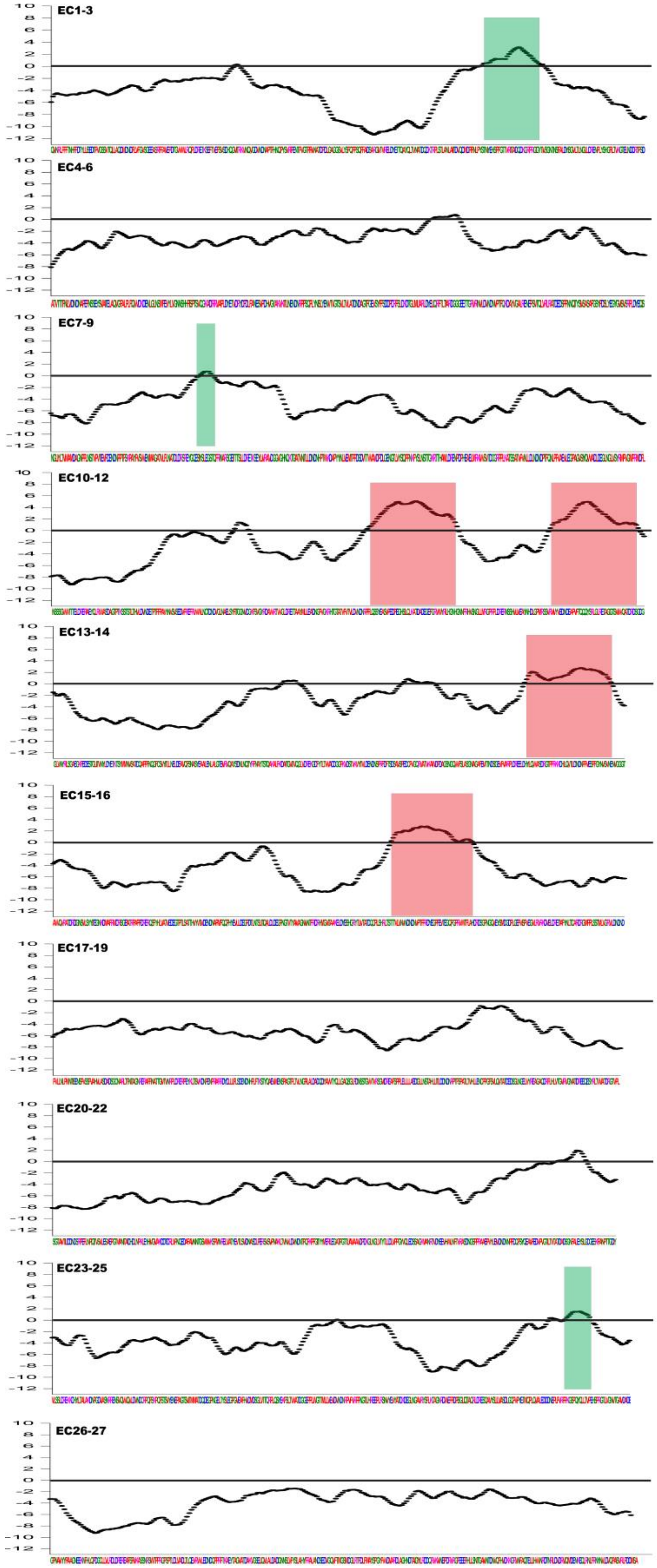
EC1-3
EC4-6
EC7-9
EC10-12
EC13-14
EC15-16
EC17-19
EC20-22
EC23-25
EC26-27
10
8
6
4
2
0
-2
-4
-6
-8
-10
-12

**Figure S1. The propensity-score of Cdh23 EC1-27 to undergo liquid-liquid phase separation with the amino-acid sequence plot**. The plot of the propensity scores (on the Y-axis) estimated using the catGRANULE algorithm for Cdh23 EC1-27 against the residues among entire EC domains (on X-axis) shows that Cdh23 has a higher probability of undergoing LLPS. EC regions with CDF scores above 1 are highlighted with red (with a high propensity for LLPS) and green (with a moderate propensity) boxes.

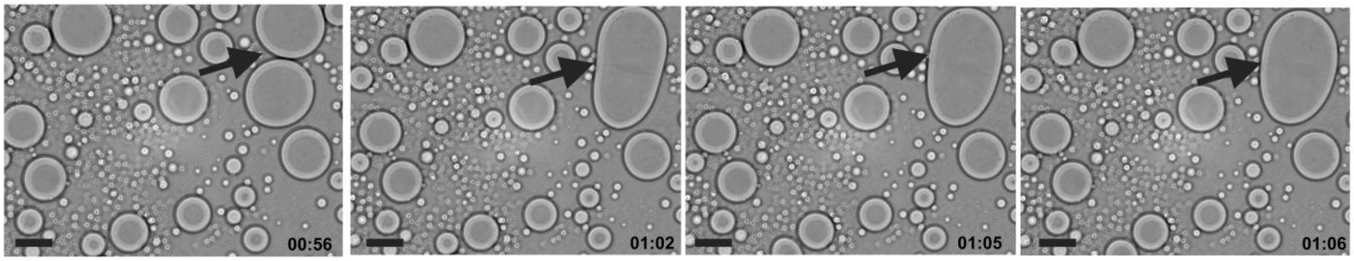


**Figure S2. Time-lapse images of droplet fusion (Supporting to Fig. 1 C).** The bright-field images with time capture one of the fusion events of liquid droplets of Cdh23 EC1-27. Arrows in black are highlighting the droplets undergoing fusion. Scale bar: 25 µm.

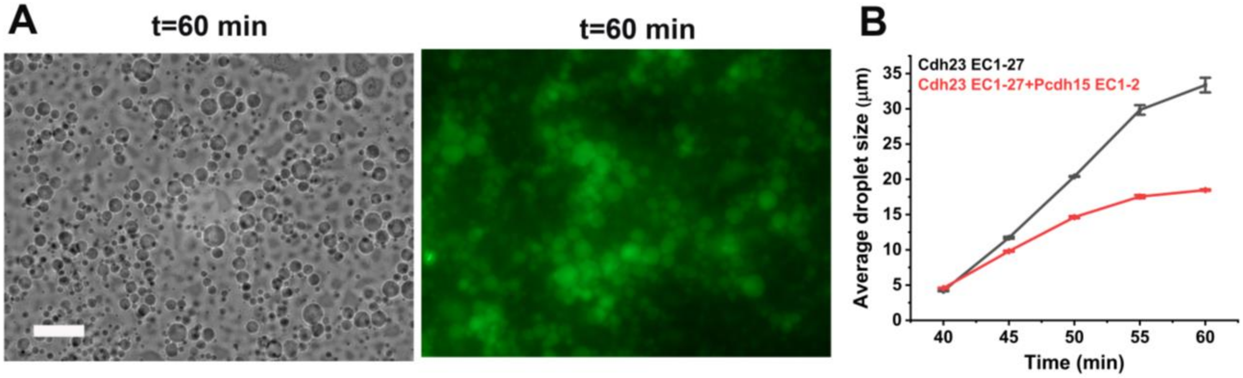


**Figure S3. LLPS of Cdh23 EC1-27 in the absence of trans-interactions. (A)** Bright-field and **(B)** fluorescence images of liquid droplets of Cdh23 EC1-27 induced by exclusive cis-interactions. The trans-interactions were turned off by introducing Pcdh15 EC1-2 in the buffer. The scale bar is 50 µm. **(C)** The comparative growth kinetics of liquid droplets (µm) of Cdh23 EC1-27 in the absence (black) and presence (red) of Pcdh15 EC1-2. Pcdh15 blocks the homophilic trans-binding interface of Cdh23.

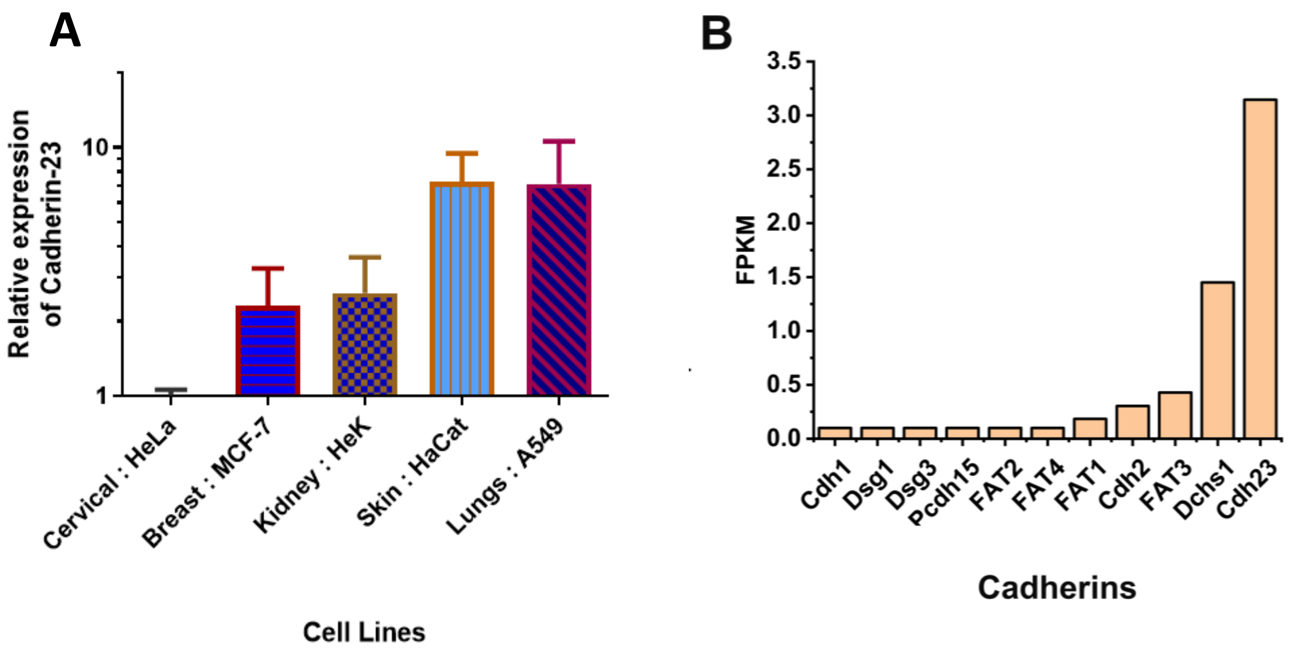


**Figure S4. Differential expression of Cdh23 in cancer cell lines and microglia (supporting to Fig. 4)**. (**A)** The relative expression of Cdh23 mRNA in different cancer cell lines, namely HeLa, MCF-7, HEK293, HaCaT, and A549, was quantified using qRT-PCR. The highest expression is noticed in A549 and the most negligible expression in HeLa. (**B)** The bar plot displays the mRNA expression of different cadherin proteins in microglia cells. FPKM is Fragments Per Kilobase Million essentially represents normalized expression values.

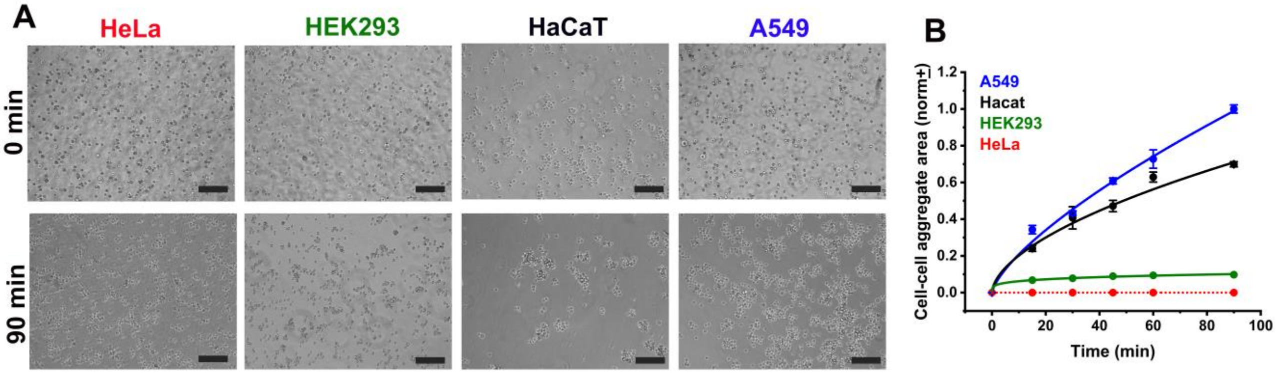


**Figure S5. Dependence of cell-cell adhesion rate on the intrinsic expression of Cdh23 (supporting to Fig. 4). (A)** Time-stamp bright-field images of cell-aggregates of HeLa, HEK293, HaCaT, and A549 cells differentially expressing endogenous Cdh23. Scale bar: 50 µm. (**B)** Growth of cell-cell aggregation area (in $\mu m^2$) with time for HeLa (red), HEK293 (green), HaCaT (black) and A549 (blue) cell lines. Error bars represent the standard error of the mean (SEM) for N=15 aggregates. The solid lines represent the fitting of aggregation kinetics to the Von Bertalanffy model. The dotted lines represent no fitting of the data.

**Video 1 A. Fusion of droplets of Cdh23 EC1-27 (Supporting to Fig. 1 C).** The fusion of liquid droplets of Cdh23 EC1-27 was captured under a fluorescence (GFP) microscope.

**Video 1 B. Fusion of droplets of Cdh23 EC1-27 (Supporting to Fig. 1 C).** The fusion of liquid droplets of Cdh23 EC1-27 was captured under a bright field.

**Table S1. Diffusion coefficients measured for the clusters of Cdh23 EC1-27 anchored to GUV membranes**

| Experimental condition | *Diffusion coefficient ($\mu m^2/s$) |
|---|---|
| Cdh23 EC1-27 labelled with nile red | 0.16 ± 0.05 |
| Cdh23 EC1-27 labelled with nile red + 1,6-HD treated | 0.114 ± 0.02 |
| Cdh23 EC1-27 labelled with nile red + Imidazole treated | 0.12 ± 0.04 |

*The diffusion of Cdh23 is calcium dependent

**Table S2. Rate constants for cell aggregation at the given conditions measured by fitting the aggregation kinetics to the Von Bertalanffy model**

| **Experimental condition** | **Rate constant ($a_{on}^{0}$) (*$min^{-1}$*)** |
|---|---|
| HEK293 | $7.96 \times 10^{-9}$ |
| HEK293 Cdh23 EC1-10 | $3.4 \times 10^{-4}$ |
| HEK293 Cdh23 EC1-27 | $6.2 \times 10^{-3}$ |
| HEK293 + 1,6-HD | $7.55 \times 10^{-9}$ |
| A549 | $7.6 \times 10^{-3}$ |
| A549 + 1,6-HD | $1.26 \times 10^{-4}$ |
| A549 + Cdh23-siRNA | $7.25 \times 10^{-4}$ |
| HaCaT | $3.4 \times 10^{-3}$ |

**Table S3. CDF scores for different cadherins (EC-domains only) estimated using the catGRANULE algorithm.**

| **Cadherins** | **CDF score** |
|---|---|
| Cdh23 | 1.259 |
| Pcdh1 | 1.002 |
| Dcsh1 | 1.350 |
| FAT1 | 1.432 |
| FAT2 | 1.326 |
| FAT3 | 1.538 |
| FAT4 | 1.642 |
